# Acute SARS-CoV-2 infection in pregnancy is associated with placental ACE-2 shedding

**DOI:** 10.1101/2021.11.19.469335

**Authors:** Elizabeth S. Taglauer, Elisha M. Wachman, Lillian Juttukonda, Timothy Klouda, Jiwon Kim, Qiong Wang, Asuka Ishiyama, David J. Hackam, Ke Yuan, Hongpeng Jia

## Abstract

Human placental tissues have variable rates of SARS-CoV-2 invasion resulting in consistently low rates of fetal transmission suggesting a unique physiologic blockade against SARS-CoV-2. Angiotensin-converting enzyme (ACE)-2, the main receptor for SARS-CoV-2, is expressed as cell surface and soluble forms regulated by a metalloprotease cleavage enzyme, ADAM17. ACE-2 is expressed in the human placenta, but the regulation of placental ACE-2 expression in relation to timing of maternal SARS-CoV-2 infection in pregnancy is not well understood. In this study, we evaluated ACE-2 expression, ADAM17 activity and serum ACE-2 abundance in a cohort of matched villous placental and maternal serum samples from Control pregnancies (SARS-CoV-2 negative, n=8) and pregnancies affected by symptomatic maternal SARS-CoV-2 infections in the 2^nd^ trimester (“2^nd^Tri COVID”, n=8) and 3rd trimester (“3^rd^Tri COVID”, n=8). In 3^rd^Tri COVID as compared to control and 2^nd^Tri-COVID villous placental tissues ACE-2 mRNA expression was remarkably elevated, however, ACE-2 protein expression was significantly decreased with a parallel increase in ADAM17 activity. Soluble ACE-2 was also significantly increased in the maternal serum from 3^rd^Tri COVID infections as compared to control and 2^nd^Tri-COVID pregnancies. These data suggest that in acute maternal SARS-CoV-2 infections, decreased placental ACE-2 protein may be the result of ACE-2 shedding. Overall, this work highlights the importance of ACE-2 for ongoing studies on SARS-CoV-2 responses at the maternal-fetal interface.

## Introduction

The COVID-19 pandemic, caused by SARS-CoV-2, has resulted in more than one hundred million infections and claimed more than 3 million lives worldwide ^1^. Despite effective vaccination and relatively rapid/accurate detection of the virus, the pandemic slowdown is hindered by frequent mutations, which led to new outbreaks in certain areas and then spread globally^2–6^. Therefore, further understanding the pathogenesis of and the host defense mechanisms to the viral infection is key in developing novel therapies to curb the spread of COVID-19.

Angiotensin-converting enzyme 2 (ACE2) is the major receptor for SARS-CoV-2 ^7, 8^. This receptor’s inducibility and variability are proposed to be an essential element for viral tropism, infectivity, and COVID-19 disease progression/outcomes^9–14^. ACE2, a vital member of the renin-angiotensin system (RAS), is a monocarboxyl peptidase and type I transmembrane protein ^15^. As a proteinase, ACE2 cleaves one amino acid from carboxyl terminals of its substrates, such as Angiotensin II (Ang II), to generate active metabolite, Ang 1-7, which counters the action of Ang II in many physiological processes, including host defense and inflammatory responses^16^. Therefore, ACE2/Ang 1-7/Mas1R axis is a negative regulatory mechanism of RAS to alleviate detrimental effects of the over-activated angiotensin-converting enzyme (ACE)/Ang II/ angiotensin receptor 1(AT1R) axis.

A majority of ACE2, after post-translational modification, migrates to the cell membrane. As a type I transmembrane protein, the anchored ACE2 contains a large ectodomain, in which the SARS-CoV-1 and −2 binding domain and enzymatic active domain separate distally, so the respective functions are not interfered with by the other ^17^. Notably, the ectodomain of ACE2 undergoes shedding constitutively, and the shedding is inducible in response to various stimuli, including bacteria and viruses. The shed or soluble ACE2 is both enzymatically active and capable of binding to existing viral particles, such as SARS-CoV-2 ^18^. The soluble ACE2 has been proposed as a decoy receptor to trap the SARS-CoV virus as a promising therapy for COVID-19, and several clinical trials are underway to test the possibility ^17^. However, controversy regarding such a strategy remains because report indicates that soluble ACE2 can facilitate viral entry via alternative routes^19^.

ACE2 shedding predominantly depends upon the activity of a disintegrin and metalloprotease domain 17 (ADAM17), or tumor necrosis factor-alpha converting enzyme (TACE) ^18, 20^. ADAM17 is one of 13 genes in the family that encode functional proteases and are involved in ectodomain shedding of an array of growth factors, cytokines, receptors, and adhesion molecules, including TNF-a and ACE-2 ^20, 21^. ADAM17 enzymatic activity is controlled by posttranscriptional tissue inhibitors of metalloprotease (TIMPs). However, its gene expression is affected by a variety of stimuli, and the sex hormonal regulation of Adam 17 has not yet been concluded ^22, 23^.

In the placenta, ACE-2 is found primarily in the outer trophoblast epithelial cell layers of the villous placenta, directly juxtaposed with maternal blood at the main functional interface between mother and fetus^24^. Placental expression levels of ACE-2 (and its partner components of the RAS) are prominent in early pregnancy and gradually decrease throughout gestation, implicating role for ACE-2 in key events of placentation ^24^. Several other groups have also identified ACE-2 expression in the villous placenta primarily in trophoblast epithelial cells with a lack of expression among other stromal cells and fetal endothelium ^25–27^. ADAM17 is also found within the villous placental trophoblast cells and its expression and its activity in the placenta has primarily been linked to TNF-a production in inflammatory pregnancy states ^28–30^.

To date, the majority of studies evaluating ACE-2 in pregnancies affected by COVID-19 have focused on maternal SARS-CoV-2 infections in the third trimester ^26, 27, 31^. As the pandemic has ensued, women have gradually experienced COVID-19 disease throughout their pregnancies with consistent reports of low SARS-CoV-2 fetal transmission. Yet ACE-2 expression relative to gestational stage of maternal SARS-CoV-2 infection remains unknown. Further, while the phenomenon of ACE-2 shedding, ADAM17 activity and serum ACE-2 levels have been characterized in multiple tissue types and COVID-19 patient populations ^32^, they are not well defined in the human placenta or in relation to the timing of maternal COVID-19 infections in pregnancy.

Given the dynamic nature of ACE-2 and ADAM17 activity among multiple tissues impacted by SARS-CoV-2 infection and COVID-19 disease evolution ^17, 32^ we hypothesized that ACE-2 expression and ADAM17 activity in the placenta is impacted by the timing of maternal SARS-CoV-2 infection in pregnancy relative to delivery. In the present study, ACE-2 expression/ADAM17 activity were evaluated in placental villous tissues and maternal serum ACE-2 levels in a cohort of maternal-fetal dyads with 2nd and 3rd trimester maternal SARS-CoV-2 infections in comparison with control pregnancies (Figure 1A). This study design allowed for analysis of ACE-2 placental expression dynamics in states of acute vs remote SARS-CoV-2 infections in pregnancy to further understand the trajectory of responses to COVID-19 at the maternal-fetal interface.

**Figure 1.**
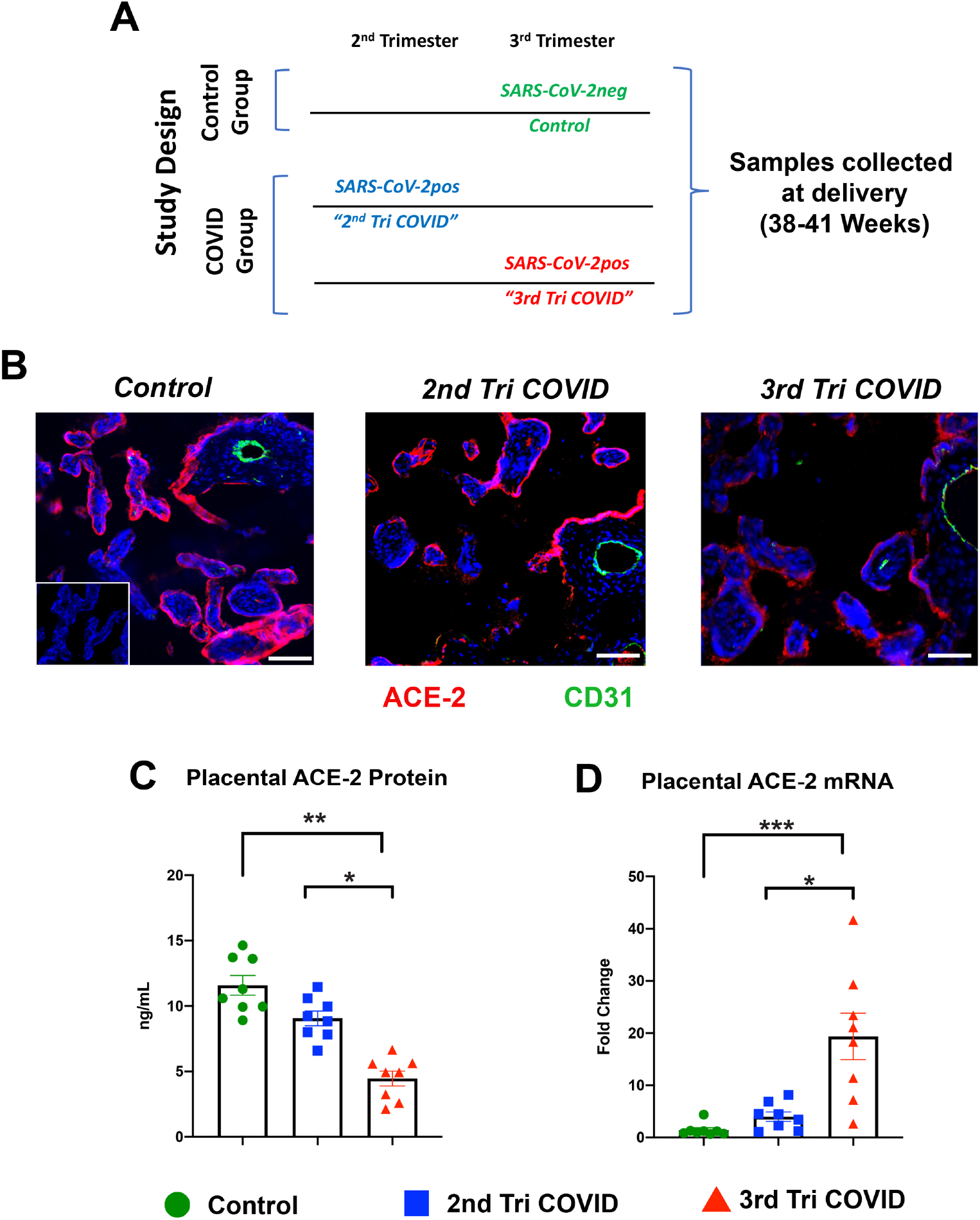
Villous placental ACE-2 expression in acute vs remote SARS-CoV-2 infections in pregnancy. **A**. Study Design. Control group: pregnancies who had no report of SARS-CoV-2 infection or COVID-19 symptoms during their pregnancy and were SARS-CoV-2 negative via universal screening at time of admission to labor and delivery. COVID group: women with documented COVID-19 symptoms and a SARS-CoV-2 positive test during their 2nd trimester (*2nd Tri COVID*) or 3rd trimester (*3rd Tri COVID*) of pregnancy. Schematic shows timing of maternal SARS-CoV-2 infection relative to sample collection at delivery. **B**. Representative images from immunohistochemical survey of ACE-2 in villous placental tissues from each patient group (n=8 per group). Red: ACE-2, Green: CD31, Blue: DAPI nuclear stain. Inset image: 2^nd^ only antibody negative control. Scale bars: 25mm. VP: Villous placenta; BV: fetal blood vessel. **C.** ACE-2 expression in villous placental tissue homogenates as assayed by Human ACE-2 ELISA. **D** qRT-PCR analysis of ACE-2 mRNA expression in villous placental tissues. Error bars: +/− standard error of the mean. * *p* < 0.05, ** *p* < 0.01 *** *p* < 0.001

## Materials and Methods

### Study enrollment

The current study was approved by the Boston University Medical School Institutional Review Board and written informed consent was obtained from all subjects. Prospective mother-infant dyads were enrolled at Boston Medical Center (BMC) between July 2020 and April 2021. Eligibility criteria were: ≥ 18 years of age, documented symptomatic SARS-CoV-2 infection during the second or third trimester of pregnancy (testing via nasal swab PCR, COVID groups) or no documented SARS-CoV-2 infection during pregnancy (testing via nasal swab PCR, Control group) singleton gestation pregnancy, English or Spanish speaking. Exclusion criteria included inability to provide informed consent.

### Sample collection and processing

Maternal blood samples were collected in EDTA collection tubes by trained staff within 24 hours pre or post-delivery. Following collection, blood samples were then stored at 4 °C and centrifuged within 6 hours of collection. Plasma was then extracted and frozen at −80°C until analysis.

For placental collection, all tissues were collected within an average time of 3.1 hours (SEM +/− 0.14) following delivery and all samples were taken midway between umbilical cord insertion site and edge of placental disk. For histological analysis, full thickness placental biopsies (1cm x 3cm) were collected and fixed in 10% neutral buffered formalin (ThermoFisher) for 72 hours, soaked in 18% sucrose for 24 hours, embedded in Tissue-Plus™ optimal cutting temperature (OCT) compound (ThermoFisher) and frozen at −80°C. For villous placental protein and RNA analysis, triplicate 0.5cm pieces of villous placental tissue (obtained from a depth midway between decidua basalis and chorionic plate) were dissected, flash frozen neat or in *RNAlater* (ThermoFisher) and stored at −80°C

### Immunohistochemistry

Fixed/frozen placental tissue blocks were cryo-sectioned at 15μm. Slides were then washed with 1X PBST(Tween20) and then blocked with donkey blocking serum for 1h. Primary antibodies of interest, including sheep anti-CD31 (1:100; AF806, R&D systems), rabbit anti-SARS-N protein (1:500; 200-401-A50 Rockland) and goat anti-ACE2 (1:50; AF933, R&D Systems) were then incubated at 4°C overnight. The next day, samples were washed five times for 10 minutes each in PBST. After blocking with donkey serum at room temperature for 1h, secondary conjugate antibodies, donkey anti-sheep Alexa Fluor 488 (1:250; 713-545-003, Jackson Immunoresearch), donkey anti-rabbit Alex Fluor 555 (1:250; 711-165-152, Jackson Immunoresearch) and donkey anti-goat Alexa fluor 647 (1:250; 705-605-147, Jackson Immunoresearch) were applied for 6h at room temperature. Slides were then washed five times with PBST again for 10 minutes each and mounted with DAPI (AF806, Vector Laboratories). Sections were imaged on a Nikon deconvolution wide-field epifluorescence microscope and processed using NIS-Elements Software (Nikon).

### Quantitative reverse transcription-polymerase chain reaction (qRT PCR) analysis

RNA was extracted from fresh frozen villous placental tissue biopsies with using RNA was isolated using an RNAqueous™ kit (Invitrogen) per manufacturer’s protocol. RNA transcripts were subsequently evaluated with Taqman^®^ probes/primers (Forward: TAATGCTGGGGACAAATGGT, Reverse: CAGCTGAAGCTTGACTGTGAG, probe: TCCACACTTGCCCAAATGTA) for ACE-2. Target expression was normalized to the transcript for GAPDH and relative expression was quantified via fold change in COVID samples relative to Control using 2^−ΔΔCT^ calculations.

### ELISA Assays

#### Villous Placental Tissue ACE-2 ELISA

Fresh frozen dissected placental tissue (300-500mg) was homogenized with protein lysis buffer (50mM Tris-HCL pH 8.0, 150mM NaCl, 1% NP-40, 0.5mM PMSF(Sigma) and cOmplete protease inhibitor (Roche). Tissue lysates were then centrifuged at 20,000 x g x 30min at 4°C followed by collection of sample supernatants. Protein concentration for each sample was determined using a Micro BCA™ Protein Assay Reagent Kit (ThermoFisher) per manufacturer’s instructions. Samples were assayed into a Human ACE-2 DuoSet ELISA assay (R&D systems), loading equal amounts of protein in duplicate for all samples.

#### Serum ACE-2 and Estradiol ELISA

Frozen maternal serum aliquots were quick thawed and immediately assayed using either a Human ACE-2 DuoSet ELISA assay (R&D systems) or a Parameter*TM* Estradiol assay per manufacturer’s instructions. All samples were assayed in duplicate. Absorbance for all ELISA assays was evaluated using a Molecular Devices SpectraMax i3x Multi-Mode Microplate Reader.

### ADAM17 activity assay

ADAM17 (TACE) activity was assayed in fresh frozen villous placental tissue homogenates using a SensoLyte(R)520 Fluorometric TACE Activity Assay kit. For sample normalization, equal amounts (300mg) of fresh frozen villous placental tissue samples were homogenized into the assay lysis buffer containing 0.1% Triton-X 100 (Sigma) and further processed per manufacturer’s instructions. Tissue homogenates were run in duplicate for each placental sample. Fluorescence intensity values (Ex/Em= 490nm/520nm) were then measured using a Molecular Devices SpectraMax i3x Multi-Mode Microplate Reader every 5 min for a total of 40 minutes.

### Statistical Analysis

Patient demographic characteristics, and maternal and neonatal outcomes were reported as follows: Continuous variables were reported as either median with interquartile ranges or means with standard deviation and categorical variables were reported as percentages as indicated for each parameter (Table 1). P-values for continuous variables were generated using a one-way analysis of variance test with Tukey’s post hoc analysis. P-values for categorical variables were generated using the Fisher exact tests. Differences were considered significant at α = 0.05. Statistical analysis for tissue and serum ACE-2 ELISA, serum estradiol ELISA and ACE-2 qRT-PCR were evaluated using a one-way analysis of variance test with Tukey’s post hoc analysis for multiple comparisons. The significance of ADAM17 activity values over time and per experimental group were evaluated using a two-way analysis of variance test with Tukey’s post hoc analysis. All statistical analyses were conducted using Prism 7 software (GraphPad, San Diego, CA).

**Table 1.** Patient Demographics.

| Demographic | Control (N=8) | 2<sup>nd</sup> Tri COVID (N=8) | 3<sup>rd</sup> Tri COVID (N=8) | P-Value |
| --- | --- | --- | --- | --- |
| Weeks gestation age at SARS-CoV-2 infection – Mean (SD) | N/A | 18.7 (4.1) | 30.0 (3.1) | N/A |
| Maternal COVID severity – N (%) |  |  |  | N/A |
| Hospitalized | N/A | 0 | 1 (12.5%) |  |
| ICU |  | 0 | 0 |  |
| Maternal age (years) – Mean (SD) | 30.4 (6.6) | 30.0 (5.0) | 30.9 (5.1) | 0.95<sup>§</sup> |
| Maternal race – N (%) |  |  |  |  |
| Black | 0 | 1 (12.5%) | 1 (12.5%) | 0.89 |
| White | 2 (25%) | 3 (37.5%) | 2 (25%) |  |
| Other | 6 (75%) | 4 (50%) | 5 (62.5%) |  |
| Maternal chronic health conditions<sup>a</sup> – N (%) | 5 (62.5%) | 3 (37.5%) | 4 (50%) | 0.87 |
| Pregnancy complications<sup>b</sup> – N (%) | 5 (62.5%) | 7 (87.5%) | 8 (100%) | 0.27 |
| Gestational age at birth (weeks) – Mean (SD) | 38.6 (0.9) | 39.9 (1.1) | 40.0 (1.9) | 0.09<sup>§</sup> |
| Infant sex – N (%) |  |  |  |  |
| Male | 3 (37.5%) | 5 (62.5%) | 2 (25%) | 1 |
| Female | 5 (62.5%) | 3 (37.5%) | 6 (75%) |  |
| Birth weight (grams) – Mean (SD) | 3353 (459) | 3453 (327) | 3446 (411) | 0.86<sup>§</sup> |
| 5-minute APGAR – median (IQR) | 9 [8.75- 9.00] | 9 [9- 9] | 9 [9 - 9] | 0.12<sup>#</sup> |
| Required NICU admission – N (%) | 1 (12.5%) | 1 (12.5%) | 0 | 1 |
| Infant SARS-CoV-2 positive results | 0 | 0 | 0 | N/A |

<sup>a</sup>Chronic health conditions included autoimmune disease, diabetes, hepatitis B, hepatitis C, HSV, HIV, hypertension, obesity, thyroid disease, substance use disorder, or other
<sup>b</sup>Pregnancy complications included chorioamnionitis, gestational diabetes, hypertensive disorder of pregnancy, fetal growth restriction, placenta previa, preterm labor, unexplained vaginal bleeding, or other.
P-values for continuous variables were generated using ANOVA test (indicated by §), or Kruskal Wallis rank test (indicated by #). P-values for categorical variables were generated using the Fisher exact tests.

## Results

To evaluate ACE-2 expression dynamics at the maternal fetal interface in varied types of SARS-CoV-2 infections during pregnancy, a cohort of matched villous placental tissues and maternal serum samples were collected at time of delivery. SARS-CoV-2 infections in pregnancy were classified according to timing relative to delivery: “remote” infections in the 2nd trimester (2^nd^ Tri COVID) or “acute” infections in the 3rd trimester (3^rd^ Tri COVID) (**Figure 1A**).

All SARS-CoV-2 positive patients (COVID group) were symptomatic at time of testing (specified as with fever or reports of respiratory or gastrointestinal symptoms and/or loss of smell/taste documented in the medical chart). All cases were classified as “mild-moderate” infections with only one case requiring hospital admission due to COVID-19 and no patients requiring respiratory support or ICU care during the symptomatic phase of their illness **(Table 1).** The SARS-CoV-2 negative cohort (Control) were tested via universal screening procedures at time of admission to BMC labor and delivery. There was no significant difference in patient race, ethnicity, mode of delivery or gestational age at delivery between groups. Additionally, infant birth outcomes (i.e. APGAR scores and birthweight) and infant gender were also not statistically different between groups. Finally, within the COVID pregnancy group, all infants who met criteria for testing were negative for SARS-CoV-2.

### Villous placental ACE-2 expression is decreased in acute maternal SARS-CoV-2 infections in pregnancy

To evaluate the abundance of ACE-2 in placental tissues from our patient cohorts, we first examined ACE-2 expression in full thickness placental biopsies using immunohistochemistry. Placental tissues were first screened for the presence of SARS-N protein. Only 3 of the 16 placental tissues within our COVID showed the presence of SARS-N expression (n=1 in 2nd trimester cohort; n=2 in 3rd trimester cohort; data not shown), a frequency similar to previously published placental tissue cohorts ^31, 33^. We first identified ACE-2 expression in villous trophoblast epithelial cells (but not fetal blood vessel endothelium), consistent with previous publications ^25–27^ **(Figure 1B).** We also noted that villous placental tissues from remote, 2^nd^ Tri COVID had similar ACE-2 expression intensity to those of Control placental tissues. However, ACE-2 expression appeared to be decreased in placentas from acute, 3^rd^ Tri COVID **(Figure 1B)** as compared to both Control and the 2^nd^ Tri COVID group.

To further quantify ACE-2 expression in these placenta tissues, we isolated protein from dissected villous placental tissue lysates and quantified ACE-2 expression through ELISA assay analysis. Similar to the trend noted in our immunohistochemical analysis, we identified that ACE-2 in acute third trimester infections was significantly decreased in comparison with Control and 2^nd^ Tri COVID groups **(Figure 1C).** Thus, through immunohistochemical and quantitative protein analysis, in our tissue cohort, ACE-2 expression in villous epithelial tissues appeared to be decreased in acute, but not remote, SARS-CoV-2 infections in pregnancy as compared with controls. Interestingly, qRT-PCR analysis of ACE-2 from villous samples of the same placental tissues showed a significant upregulation of ACE-2 mRNA expression in acute 3^rd^ Tri COVID (**Figure 1D**), suggesting the noted decrease placental ACE-2 protein was due to post-translational influences rather than alteration of mRNA transcripts.

### Acute maternal SARS-CoV-2 infections in the third trimester are associated with increases in placental ADAM17 activity and circulating maternal serum ACE-2

To next investigate the potential post-translational mechanism behind decreased placental ACE-2 expression associated with acute maternal SARS-CoV-2 infections, ADAM17 activity was evaluated in dissected villous placental tissues. As a cell surface enzyme known to cleave surface ACE-2 ^20^, we examined whether the activity of ADAM17 in the placenta varied in relation to the timing of maternal SARS-CoV-2 infections in pregnancy. For this analysis, we used a fluorescence-based assay evaluating kinetic ADAM17 cleavage activity over time ^34, 35^. We first noted that in comparison with our control group, placental tissues from remote SARS-CoV-2 infections in the 2nd trimester of pregnancy showed a significant difference in ADAM17 activity (**Figure 2A).** However, placental tissues from pregnancies with acute third trimester maternal SARS-CoV-2 infections had a significant increase in ADAM17 activity over time when compared with control villous placental tissues (**Figure 2B**).

**Figure 2.**
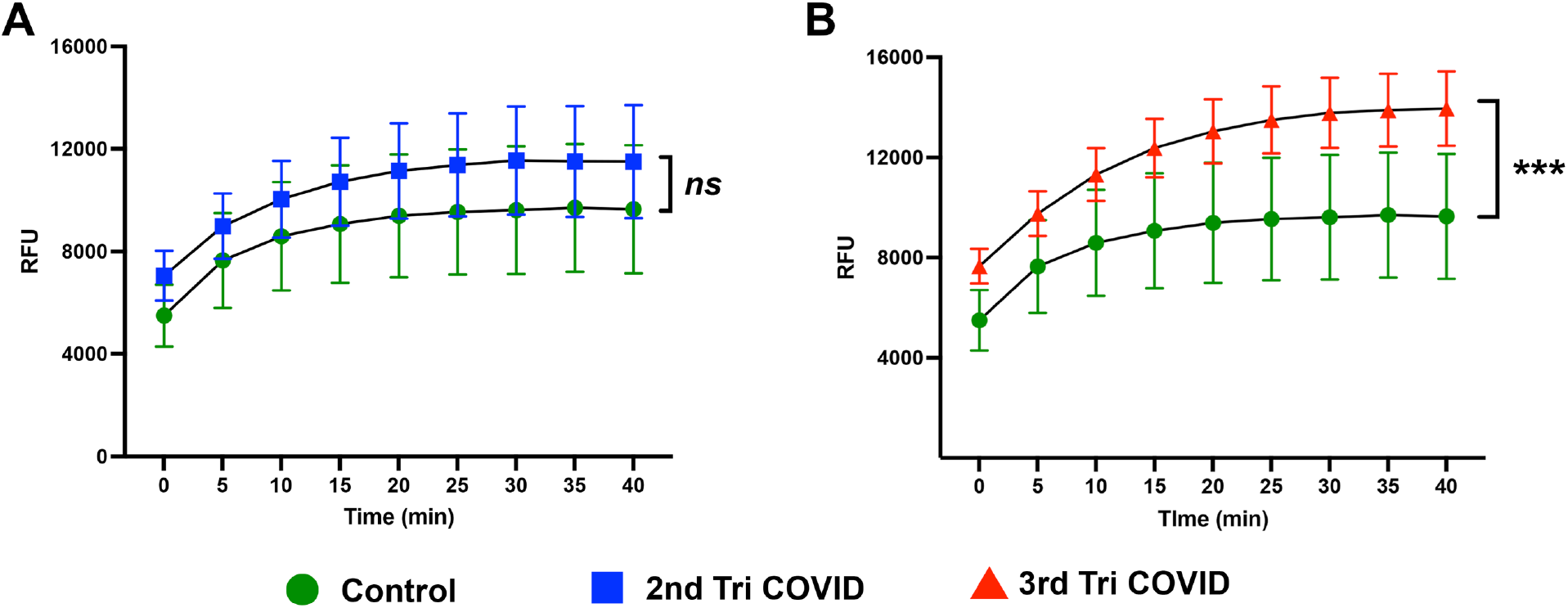
Increased placental ADAM17 activity in acute maternal SARS-CoV-2 infections during pregnancy. ADAM17 activity over time in villous placental tissue homogenates in placental tissues from patient groups as described in Figure 1. A. Control group vs 2nd Tri COVID. B. Control group vs 3rd Tri COVID. Error bars: Standard error of the mean; RFU: Relative fluorescent units. \*\*\**p* = 0.003.

The combined results of decreased villous placental ACE-2 expression, increased *ACE-2* gene expression, and increased ADAM17 activity suggested a cleavage of ACE-2 from the villous placental tissue. As villous placental tissue is directly juxtaposed with the maternal blood, we next examined whether the changes we observed in ACE-2 expression and ADAM17 activity also correlated with any alterations in circulating ACE-2 levels in maternal serum. To conduct this analysis, we evaluated ACE-2 abundance in maternal serum samples which matched with placental samples analyzed for ACE-2 expression and TACE activity. In line with the trends noted with ADAM17 activity, serum ACE-2 was significantly increased in acute 3rd trimester SARS-CoV-2 infections as compared to control and 2nd trimester maternal SARS-CoV-2 infections (**Figure 3A)**.

**Figure 3.**
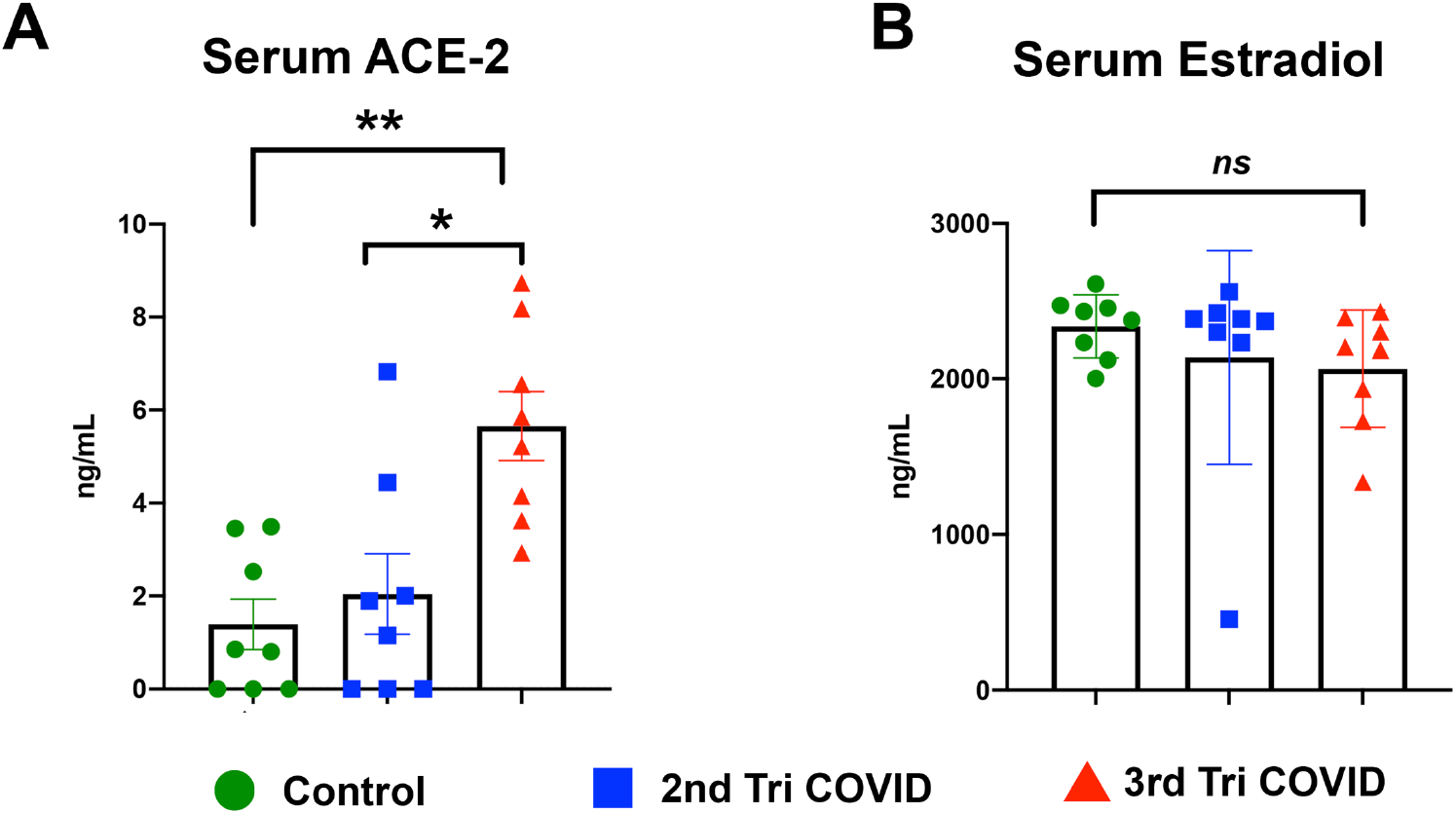
Maternal SARS-CoV-2 infection in the third trimester is associated with increased circulating maternal serum ACE-2. Soluble ACE-2 (A) and serum estradiol (B) in maternal serum collected at delivery from patient groups as described in Figure 1. Error bars: standard error of the mean. \*\**p*= 0.0063; \**p* = 0.043.

Finally, we evaluated whether changes in placental and serum ACE-2 abundance correlated with alterations in circulating estrogen levels in our patient cohort. Estogen regulates ACE-2 expression within airway epithelial cells ^36^ and has been implicated in the gender differences noted between COVID-19 severity among adults ^37^. However, the role of estrogen on ACE-2 regulation in pregnancy has been less well defined. Among our cohort there was no significant difference in maternal serum estrogen (assayed as estradiol) levels between our control and COVID groups (**Figure 3B**).

## Discussion

The current study identifies new evidence on the dynamics of placental ACE-2 expression in SARS-CoV-2 infections in pregnancy and the influence of the gestational stage of maternal infection relative to delivery. Our findings suggest that the ACE-2 protein expression changes are the result of placental ACE-2 shedding mediated by ADAM17 in response to maternal SARS-CoV-2.

The dynamic nature of placental ACE-2 expression that we have noted in our study could account for the variation of reports identifying the presence or absence of ACE-2 within trophoblast epithelial cells ^38^. Subsequent studies ^25–27^ have strongly substantiated pre-pandemic publications ^24^ for the presence of ACE-2 the maternal fetal interface. The 3^rd^-Tri COVID-specific increase in ACE-2 mRNA could be due to compensatory upregulation of ACE-2 transcripts in response to the active shedding process in this disease state. Recently, a truncated form of human ACE-2, namely delta ACE-2 has been reported to be upregulated by interferons^39^. Our primer/probe set flanks only the region in the prototype ACE-2, therefore the expression we detected is exclusive of delta ACE-2. Additionally, there were no significant differences in the maternal serum estradiol between our patient groups, further suggesting influences other than hormonal signaling affecting the dynamic expression of placental ACE-2 in these pregnancies. Ongoing evaluation of other influences on ACE-2 post-translational regulation and ADAM17 activity will be required in future studies to characterize the mechanisms governing this process more clearly. Given the tissue inflammation noted in placentas from pregnancies affected by acute maternal SARS-CoV-2 ^25, 33, 40^, pro-inflammatory cytokine regulation and immune response signaling pathways will be a key area of focus for future analysis of ACE-2 and ADAM17 regulation in these tissues.

Consistent with previous studies ^25–27^ we also found that ACE-2 was not expressed endothelium of fetal blood vessels in Control or COVID placental tissues. These findings highlight an important contrast in the pulmonary and placental responses to SARS-CoV-2. In healthy lung tissue, the pulmonary endothelium also has minimal expression of ACE-2 but in patients with fatal COVID-19, ACE-2 expressions is increased on the pulmonary endothelium, and SARS-CoV-2 invasion into primary endothelial cells is dependent on ACE-2 induction, a process facilitated by Type I interferon alfa and - beta *in vitro*^41^. In stark contrast with the lung, the lack of ACE-2 on the fetal endothelium in COVID placental tissues supports the likely central role of ACE-2 as gatekeeper protecting against perinatal transmission in pregnancies affected by maternal COVID-19.

This work is also the first study to evaluate serum ACE-2 levels in pregnant women affected by COVID-19. Serum ACE-2 levels are increased in pregnancy relative to non-pregnant women and further relative increases have been associated with small for gestational age (SGA) fetal growth ^42^. While serum ACE-2 in non-pregnant adult COVID-19 patients has been widely evaluated ^43–45^, no studies to date have characterized maternal serum ACE-2 content in pregnancies with maternal SARS-CoV-2 infection. While our combined data suggest that the increase in maternal serum ACE-2 in acute 3^rd^ Tri COVID pregnancies could be the result of increased ACE-2 placental shedding, the source of circulating maternal serum ACE-2 levels in these pregnancies (lung, placenta or both) requires ongoing investigation.

This study has several limitations. First, the cohort is a small sample size and the gestational evaluation only spanned infections in the 2nd and 3^rd^ (but not 1^st^) trimesters of pregnancy. Additionally, as stated above, soluble ACE-2 in maternal serum cannot be directly identified as placental in origin and could be derived from multiple sources. Ongoing mechanistic studies, including relevant animal models, are needed to more completely evaluate ACE-2 expression relative to maternal COVID-19 infection in all trimesters of pregnancy, to clarify the placental origin of maternal serum ACE-2 and to functionally correlate ADAM 17 activity with ACE-2 placental regulation.

In conclusion, our work highlights a previously unrecognized dynamic expression of ACE-2 in the human placenta which can impacted by the timing of maternal SARS-CoV-2 infection in pregnancy relative to delivery. Taken together our data provide support for the growing body of evidence on the importance of ACE-2 regulation in the placental response against SARS-CoV-2. The human placenta has many functional and structural parallels with the human lung ^46^, and thus continues to be an important primary human tissue for identification of key targets to combat COVID-19 pathogenesis.

## Acknowledgements

We would like to thank the Department of Pathology at Boston Medical Center (BMC); the Boston University (BU) School of Medicine Cellular Imaging Core, the BU Analytical Instrumentation Core and the Maxwell Finland Laboratory for Pediatric Infectious Disease at BMC, particularly lab manager Yazdan Dasthagrasaheb, and laboratory technician Loc Truong. We would also like to thank the staff on the BMC Labor and Delivery and Postpartum Units, as well as the nursing and physician staff of the nursery and NICU at BMC who assisted with maternal blood sampling and subject recruitment.

## Notes

***Funding Sources***: This work was funded by the Boston University Clinical and Translational Science Institute COVID-19 Pilot Grant Program, UL1TR001430 (ET,EW); This work is also supported by research grants from the National Institutes of Health (R21AI14932, and R01AI148446 to H.J).

### Competing Interest Statement

The authors have declared no competing interest.

